# Leveraging joint mechanics simplifies the neural control of movement

**DOI:** 10.1101/2021.03.02.433588

**Authors:** Daniel Ludvig, Mariah W. Whitmore, Eric J. Perreault

**Affiliations:** Department of Biomedical Engineering, Northwestern University; Shirley Ryan AbilityLab; Department of Physical Medicine and Rehabilitation, Northwestern University

**Keywords:** joint mechanics, motor control, muscle activation, perceived difficulty

## Abstract

Behaviors we perform every day, such as manipulating an object or walking, require precise control of interaction forces between our bodies and the environment. These forces are generated by active muscle contractions, specified by the nervous system, and by joint mechanics, determined by the intrinsic properties of the musculoskeletal system. Depending on behavioral goals, joint mechanics might simplify or complicate control of movement by the nervous system. However, whether humans can exploit joint mechanics to simplify neural control remains unclear. Here we evaluated if leveraging joint mechanics can simplify neural control by comparing performance in three tasks that required subjects to generate specified torques about the ankle during imposed sinusoidal movements; only one task required torques that could be generated by leveraging the intrinsic mechanics of the joint. We developed a novel approach that used continuous estimates of impedance, a quantitative description of joint mechanics, and measures of muscle activity to determine the mechanical and neural contributions to the net ankle torque generated in each motor task. We found that the torque resulting from changes in neural control was reduced when ankle impedance was consistent with the task being performed, resulting in a task that required less muscular effort. Subjects perceived this task to be easier than those that were not consistent with the impedance of the ankle and were able to perform it with the highest level of consistency. These results demonstrate that leveraging the mechanical properties of a joint can simplify task completion and improve performance.

**KEY POINTS:**

- Interacting with our environment requires production of interaction forces, which are generated by muscle contractions, specified by the nervous system, and by joint mechanics, determined by the intrinsic properties of the musculoskeletal system.
- We assessed whether leveraging joint mechanics can simplify neural control by having subjects complete 3 tasks, only one of which could be accomplished by leveraging joint mechanics.
- We found that subjects reduced their muscular effort, perceived the task to be easier and completed the task more consistently when the mechanics of the ankle were consistent with the task.
- These results highlight the importance of considering limb mechanics when interpreting measures of neural control related to movement and may benefit the design of mechanical interfaces that optimize human performance.

## Introduction

Completing motor tasks that require contact is dependent on an ability to regulate the relationship between limb motions and interaction forces with the environment. The nature of this relationship depends on the requirements of each task. For example, hopping requires joint torques that increase sufficiently upon impact so as to launch the hopper into the air (Farris & Sawicki, 2012), whereas landing from a jump requires a decrease in joint torques after the initial impact to cease motion and stabilize the body (Decker *et al*., 2003). The flexibility to perform these similar but contrasting actions arises from our ability to coordinate joint motions and torques across a range of functionally relevant situations.

Two strategies for coordinating limb motions and interaction forces are leveraging the mechanical properties of the limb associated with the current state of the neuromuscular system, and actively regulating joint torques or motions by changing the state of the neuromuscular system as can occur through changes in muscle activation. The mechanical properties of a limb or joint are often quantified by estimates of impedance, the dynamic relationship between imposed motions and the resulting torques (Hogan, 1985b). By setting impedance to a desired value, it is possible to achieve a variety of motion-torque relationships, though these are of course limited to relationships that are physiologically plausible. For example, the static component of limb impedance—often referred to as stiffness—serves to stabilize the limb by generating torques that oppose externally implied motions and increase with increasing rotation of a joint. Therefore, impedance control might be sufficient for hopping where joint torques increases with increasing joint excursion (Farris & Sawicki, 2012). However, for landing from a jump in which joint torques tend to decrease—following an initial increase—with increasing joint excursion (Decker *et al*., 2003), an impedance control strategy alone would not be possible. The alternative is to change the state the state of the limb continuously throughout the task so as to achieve the desired motion-torque relationship; this is the only feasible solution when the impedance established by the current state of the neuromuscular system is not sufficient for the demands of the task being performed.

There are many conditions in which impedance regulation provides a simple and effective control strategy for stabilizing limb posture or movement trajectories. Impedance has been shown to be regulated in many postural tasks (Finley *et al*., 2012; Krutky *et al*., 2013; Zenzeri *et al*., 2014), often to stabilize the human limb against unpredictable disturbances. These adaptations can occur through changes in limb configuration (Mussa-Ivaldi *et al*., 1985; Tsuji *et al*., 1995; Trumbower *et al*., 2009; Krutky *et al*., 2013), volitional muscle activation (Hogan, 1984; De Serres & Milner, 1991; Mirbagheri *et al*., 2000), or involuntary activation through reflex pathways (Sinkjaer *et al*., 1988; Doemges & Rack, 1992; Mirbagheri *et al*., 2000; van der Helm *et al*., 2002). The same mechanisms can be used to stabilize a limb during movement (Burdet *et al*., 2001; Franklin *et al*., 2007; Ludvig *et al*., 2017), but it is unclear if humans leverage impedance to regulate the movement-torque characteristics necessary to complete a task in which stability is not the primary concern. There is evidence from animal literature suggesting that various animals leverage the intrinsic impedance of their limbs to complete different tasks (Dickinson *et al*., 2000). For example, turkeys and wallabies rely on the impedance of the passive structures within the muscle-tendon unit to regulate motion-torque characteristics during running (Roberts *et al*., 1997) and hopping (Biewener *et al*., 1998), respectively. However, it remains unknown whether leveraging the limb mechanics in these cases simplifies the neural control of the movement.

It is necessary to actively regulate joint torques through changes in muscle activation when task demands are not compatible with impedance control. This approach has the advantage of being flexible enough to handle all movement-torque profiles that are possible to learn. However, such a strategy may require more complex neural control and associated decreases in performance or increases in cognitive demand relative to an impedance control strategy. These complexities may arise from external factors such as the unpredictable mechanical properties of the environment (Johansson & Westling, 1988), or internal factors such as the nonlinear length-tension (Gordon *et al*., 1966) and force-velocity (Wickiewicz *et al*., 1984) properties of muscle or the inherent noisiness of muscle activation (Carlton *et al*., 1985; Jones *et al*., 2002; Tracy *et al*., 2005). Currently, it is unclear if humans employ a consistent control strategy of regulated muscle activity or if they switch strategies to leverage the impedance of a limb when it is consistent with task demands.

The purpose of this study was to determine if humans leverage the impedance of a limb to complete a motor task more simply and efficiently when that impedance is aligned with task demands. We evaluated this by determining how the strategy chosen by the subject influenced the perception of difficulty and task performance. All experiments were performed on the human ankle. Subjects were required to complete three tasks differing in the required coordination between ankle motions and ankle torques. One of the tasks had a motion-torque profile consistent with the physiological impedance of the ankle, allowing it to be completed either by leveraging the impedance of the ankle or actively regulating ankle torques through changes in muscle activation. The other two tasks could only be completed by explicitly regulating ankle torques through changes in muscle activation, allowing us to evaluate the influence of this strategy on task performance. We computed ankle impedance continuously throughout the experiment while simultaneously measuring the activity of the major muscles crossing the ankle. These measures were used to determine the contributions of ankle impedance and changes in neural control to the net ankle torque. We hypothesized that there would be a decrease in ankle torque due to changes in muscle activity when impedance control was a feasible option, suggestive of a simplified neural control strategy. Our objective measures were compared to subject perceptions on the difficulty of each task. These results clarify the conditions in which impedance control is used and demonstrate the impact of that use on task difficulty and performance.

Portions of this work have been previously presented in abstract form (Ludvig *et al*., 2020).

## METHODS

### A. Ethical Approval

Twenty unimpaired adults (7 female, 13 male; 27 ± 3 years) participated in this study. All subjects provided informed consent to the protocol, which was approved by the Northwestern University Institutional Review Board.

### B. Apparatus

We secured each subject’s right ankle to an electric rotary motor (BSM90N-3150AF, Baldor, Fort Smith, AR) via a custom fiberglass cast (Fig. 1). The cast encased the entire foot but did not cover the ankle joint, preserving full range-of-motion. We aligned the ankle to the center of rotation of the motor and restricted movement to the sagittal plane. Subjects sat reclined with their hips at 135 deg and their right leg extended in front of them. We fixed the right knee at 15 deg of flexion using a brace (Innovator DLX, Össur, Reykjavik, Iceland) and secured it, along with the torso, to the chair using straps. We recorded the ankle angle using an encoder integrated with the motor. We used a 6-degree-of-freedom load cell (45E15A4, JR3, Woodland, CA) to acquire force and torque data about the ankle. We controlled the motor using a position control scheme, so the position of the subject’s ankle was always dictated by the position of the motor.

**Figure 1.**
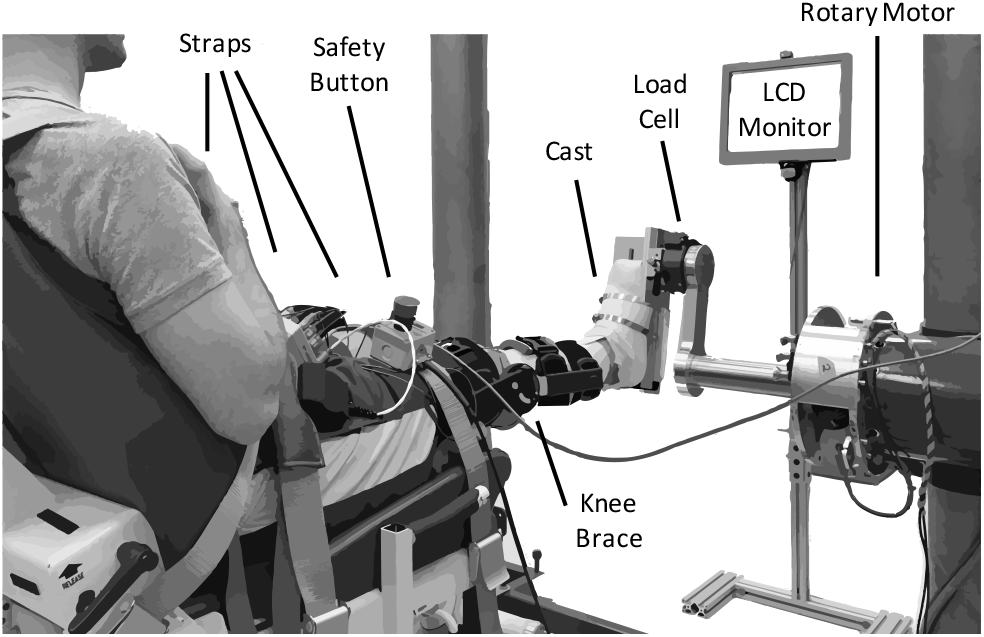
Experimental apparatus.

We measured electromyograms (EMGs) from four muscles crossing the ankle — medial and lateral gastrocnemius (MG and LG), soleus (SOL), and tibialis anterior (TA). Measurements were made using bipolar surface electrodes (Noraxon 272, Scottsdale, AZ), and amplified (AMT-8, Bortec, Calgary, AB) as needed to maximize the range of the data acquisition system. The analog data were anti-alias filtered at 500 Hz using a 5-pole Bessel filter and sampled at 2.5 kHz (PCI-DAS1602/16, Measurement Computing, Norton, MA). Ankle position was simultaneously recorded using a 24-bit quadrature encoder card (PCI-QUAD04, Measurement Computing, Norton, MA). Data acquisition and motor control were executed using xPC target (The Mathworks Inc., Natick, MA).

### C. Protocol

Two different experimental sessions were conducted. A unique set of 10 subjects participated in each session. The goal of the first session was to characterize the contribution of impedance and muscle activation to torque generation during movement. The goal of the second session was to evaluate the perceived difficulty of the three tasks performed in both sessions.

The first session began with the collection of maximum voluntary contractions (MVCs) to normalize the recorded EMG and to scale the torque for the later trials (Besomi *et al*., 2019). A MVC was collected for plantarflexion and dorsiflexion, with the ankle fixed at the neutral posture, set to be 1.75 rad between the shank and the foot. We defined ankle angle to be positive when dorsiflexed from the neutral position, consistent with previous work that has quantified ankle impedance (Mirbagheri *et al*., 2000). Since the goal of the experiment was to produce torques in the plantarflexion direction, we defined plantarflexion torque to be positive.

Subjects completed three tasks: 1) a positive-stiffness task (+K); 2) a zero-stiffness task (0K); 3) and a negative-stiffness task (-K). During all tasks, the actuator moved the ankle through a sinusoidal motion with a frequency of 0.5 Hz and an amplitude of 0.18 rad, centered about the neutral position (Fig. 2A). This frequency and amplitude were selected as they are similar to the ankle kinematics during walking (Borghese *et al*., 1996). For the +K and -K tasks, subjects were instructed to produce a 0.5-Hz sinusoidal plantarflexion torque ranging from 0–30% MVC and were aided by visual feedback. The magnitude of the target torque was selected to be feasible without fatigue over the duration of our experiments. For the +K task, the desired torque was in phase with the movement, while for the -K task the desired torque was 180° out of phase with the movement. For the 0K task, subjects were instructed to maintain plantarflexion torque constant at 15%. The +K task resulted in an angle-torque relationship with a positive slope (Fig. 2B) and hence an effective stiffness that was consistent with the impedance of the ankle. In contrast, the 0K and –K tasks had zero and negative slopes respectively, and could not be achieved simply by altering the mechanical impedance of the ankle. In all tasks, subjects were provided visual feedback of their torque and the target torque trajectory. Subjects were allowed to practice each task until proficiency was reached. We subsequently collected five 150-s trials for each of the three tasks. An additional trial was collected to determine the passive mechanics of the ankle. This involved applying the same sinusoidal movement to the ankle while subjects remained relaxed. A small pseudo-random binary sequence (PRBS) perturbation was imposed on the larger sinusoidal movement to estimate ankle impedance. The PRBS perturbation had an amplitude of 0.035 rad, a velocity of 1.75 rad/sec, and a switching time of 0.153 sec (Fig. 3). Finally, each subject tracked the sinusoidally varying target torque while the ankle position was held constant. The data from this isometric trial was used to estimate the relationship between changes in muscle activation and ankle torque.

**Figure 2.**
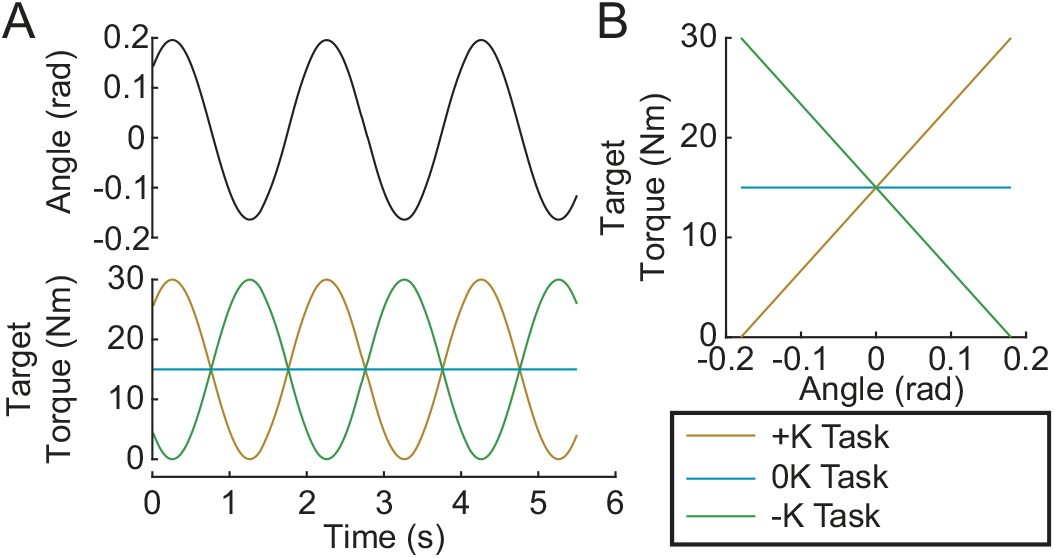
Experimental Protocol. Subjects performed 3 tasks: a positive stiffness task (+K); a zero stiffness task (0K) and negative stiffness task (-K). A) In the +K task the target torque was in phase with the imposed sinusoidal ankle rotation, while it was out of phase for the -K task. B) In the +K task, the slope of the target torque-angle trace is positive, thus we termed it a positive stiffness task. Similarly, the slope of the target torque-angle trace is zero in the zero stiffness (0K) task, and negative in the negative stiffness (-K) task.

**Figure 3.**
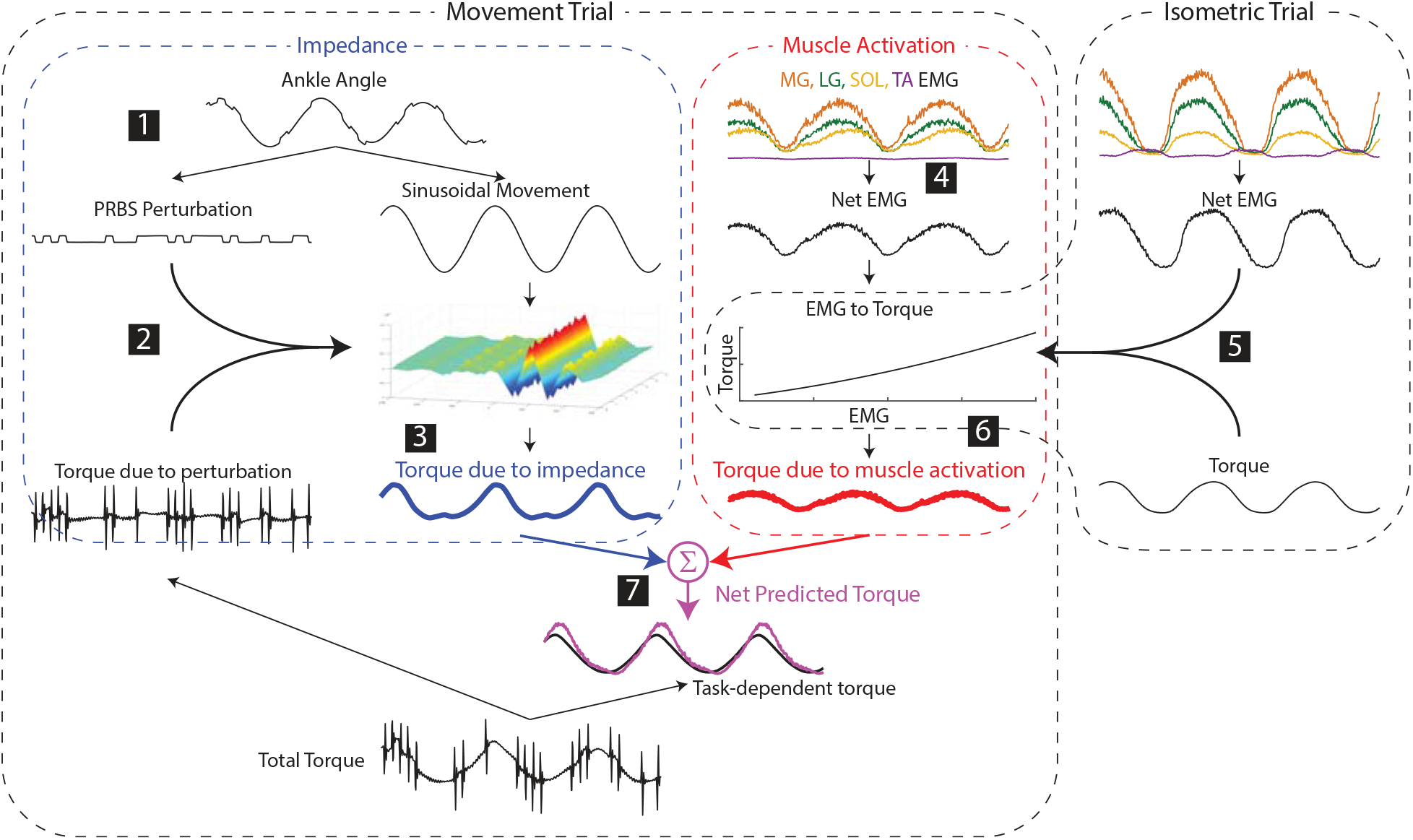
Estimating the contributions of impedance and muscle activation to the net torque about the ankle. 1) Small pseudorandom binary sequence perturbations (PRBS) were superimposed on the larger sinusoidal movement. 2) Ankle impedance was estimated by fitting a time-varying impulse response function between the small perturbation and the resultant torque. 3) The torque due to the impedance was computed by convolving the larger sinusoidal movement with the estimated impedance. 4) The net muscle activity about the ankle was approximated as the difference between the average of the plantarflexor activity and the dorsiflexor activity. 5) The torque due to muscle activation was predicted by a static relationship between the net EMG and ankle torque, estimated from data collected in a separate isometric experiment for each subject. 6) The torque due to muscle activation in the movement trials was computed using this static relationship and the EMGs measured in the movement trial. 7) The net predicted torque was computed as the sum of the torques due to impedance and muscle activation.

Task difficulty was assessed in a separate group of subjects who completed the same three tasks as in the first session, but without superimposed perturbations. Subjects completed one 60-s trial for each task, and the order of the trials was randomized. After completing all trials, subjects assigned a difficulty score to each task using a continuous scale from 1–5, with 1 defined as very easy and 5 as very hard.

### D. Estimating contributions of impedance and muscle activation to ankle torque

Prior to analysis, the recorded EMGs were notch-filtered to remove 60-Hz noise and full-wave rectified. The angle, torque and rectified EMG signals were digitally filtered to prevent aliasing and decimated to 100 Hz. Ankle impedance was estimated using an ensemble system identification algorithm that requires numerous replications of a repeatable behavior (Ludvig & Perreault, 2012). We therefore segmented all signals into overlapping three-period long segments, with each segment beginning one period (2 s) after the previous one. This resulted in approximately 370 segments for each task. We used the 200 segments with the lowest mean-squared error between measured torque and the target torque to maximize the similarity of our repetitions used for system identification. Finally, torque and EMG were normalized by each subject’s MVC torque/EMG to facilitate comparisons across subjects.

### 1) Estimation of impedance contributions to torque

Following this pre-processing, the torque due to impedance was computed as follows (numbers correspond to the steps shown in Figure 3):

1. The large sinusoidal movement and small random perturbations—as well as the torque due to the sinusoidal movement and the random perturbations—were separated by removing the ensemble mean of the ankle angle and torque from each periodic segment.
2. Ankle impedance was estimated by computing a nonparametric, time-varying impulse response function (IRF) at each point within the periodic ankle motion (Ludvig & Perreault, 2012). This impulse response described the relationship between the small PRBS perturbations and the ankle torques opposing them. Two-sided IRFs were estimated with a duration ranging from -0.06 to 0.06 s relative each instance in time; the estimation used a 0.1-s window of data centered about this same time point. (the mean %VAF of the time-varying IRFs in all subjects and tasks was 81 ± 10%, n = 30). The nonparametric IRFs were subsequently parameterized using a second order model consisting of a stiffness, viscosity and inertia (Ludvig *et al*., 2011; Ludvig & Perreault, 2012).
3. The torque due to the impedance was computed by convolving the estimated time-varying IRF with the imposed sinusoidal movement.

This procedure was done for all three tasks, as well as the data collected in the passive trial.

### 2) Estimation of muscle activation contributions to torque

Following the initial EMG pre-processing outlined above, the torque due to muscle activation was computed as follows:

4 For all tasks, the net EMG was computed by computing the difference between the average plantarflexor (LG, MG, SOL) EMG activity and the dorsiflexor (TA) EMG activity.
5 A 2^nd^ order polynomial was fit between the net EMG and the torque measured in the isometric task (%VAF = 98.0 ± 0.8%, n = 10) to create a model of the EMG to torque relationship.
6 The torque due to muscle activity during the movement trials was predicted from the EMGs measured in these trials and the isometric model.
7 The net predicted torque was computed by summing the torque due to muscle activity with the torque due to impedance computed in step 3. Note that these model-based predictions were not optimized to match the net experimental torques measured in any of the movement trials.

### 3) Evaluation of Task Performance

Task performance was evaluated by how well subjects matched the target torque, how consistent they were from trial to trial, and whether they had any consistent deviations from the target. Overall performance was quantified by the total error, quantified by the RMS of the total tracking error. Consistency was quantified by the RMS of the trial-to-trial variability, a measure of random error. Finally, the bias error, computed as the RMS of the average error, was used to quantify consistent deviations from the target torque.

### E. Statistical Analysis

The goal of this study was to determine how leveraging joint impedance when feasible simplified the neural control of movement. Specifically, we tested the hypothesis that tasks that leveraged the impedance of the ankle would be completed more simply and efficiently. We compared three metrics across the three tasks that were studied: the torque due muscle activation, perceived difficulty across the tasks, and performance in each of the three torque-tracking tasks. We used a repeated measures ANOVA to test for differences in each of these metrics across the three tasks. Post hoc analyses were computed using Tukey’s Honest Significant Difference when needed. Additionally, we ran paired t-tests to determine whether impedance or muscle activation was greater in each task. For all tests, significance was set to p = 0.05. Averages of each metric are presented as mean ± standard deviation, with accompanying number of samples. Differences between tasks are presented as mean and 95% confidence intervals (%95CI). We completed the data analysis in MATLAB (2017a, MathWorks).

## RESULTS

### A. Separating net torque into contributions from impedance and muscle activation

We found that the experimentally measured ankle torque was modeled well by our simple model predicting the torques due to impedance and muscle activation. Separating the measured torque into these two components allowed us to investigate how these two potential mechanisms for regulating motion-torque coordination were employed in each of the tested tasks. Figure 4A shows the experimentally measured torque, the predicted torques due to impedance and muscle activation, and the net predicted torque (impedance + muscle activation) for a typical subject. Across all movement trials, the standard deviation of the residual error of this model was 4.0 ± 1.6% MVC (mean ± S.D.; n = 30). This was a rather small error relative to the large torques produced in certain movement trials that were up to 30% MVC.

**Figure 4.**
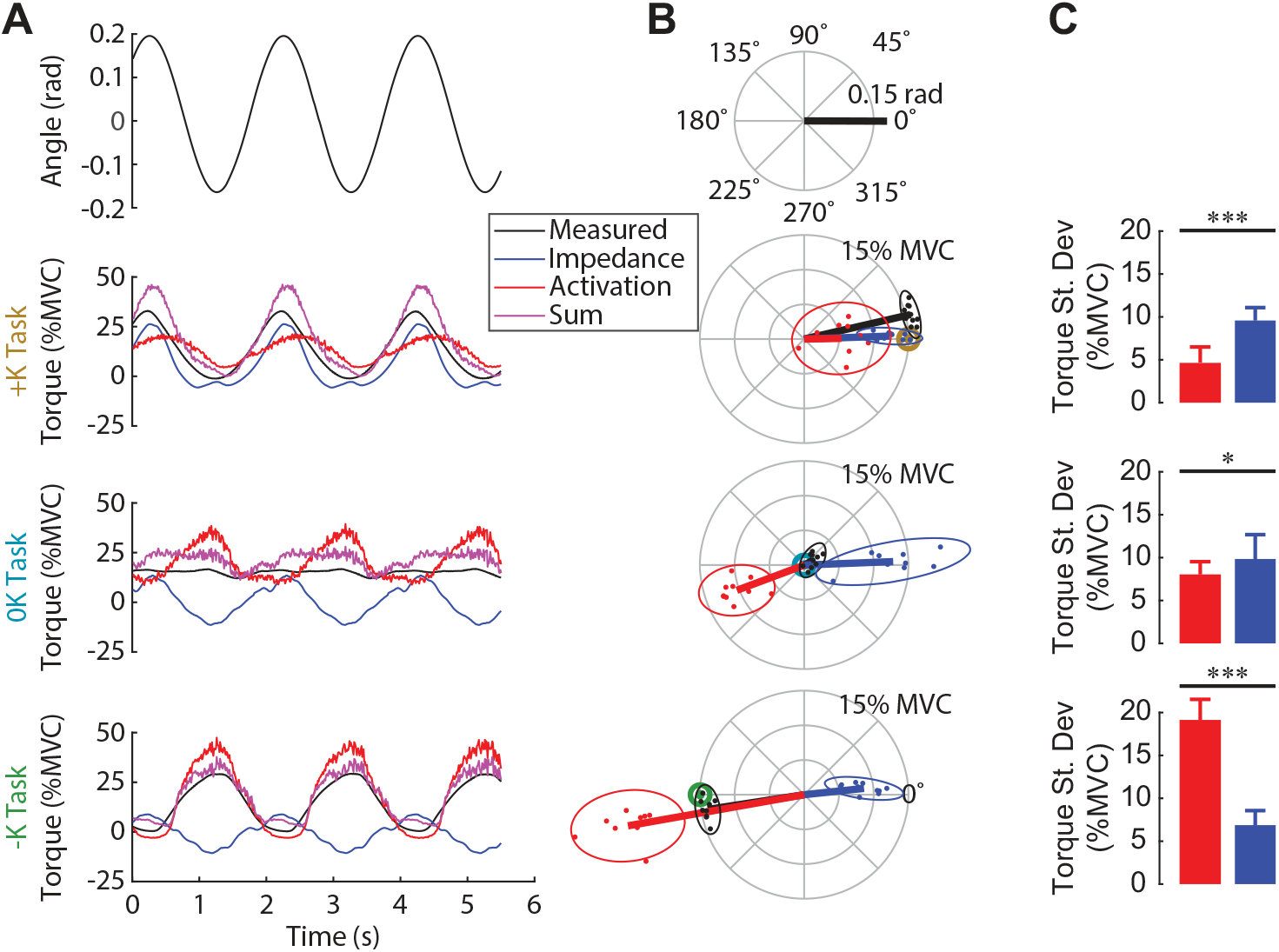
Contribution of impedance and muscle activation to the torque generated in the three tasks. A) Ankle angle, measured torque, torque due to impedance, torque due to muscle activation and their sum for all three tasks for one subject. The sum of the two modeled torque components was a good fit for the measured torque for this subject in all 3 tasks. B) Polar plot showing the phase and magnitude of the different torque components for all subjects. For each task, the target torque is denoted by a bullseye. Since the torque due to impedance was always in phase with the imposed movement, muscle activation was required to compensate for the impedance when it was not beneficial to task performance. C) The torque due to impedance was greater than the torque due to muscle activation in the +K task, whereas these torques were of similar magnitude in the 0K task, and had a reversed order of dominance in the – K task (*: p <.05, **: p < .01; ***: p < .001). We conclude that subjects completed the +K task by relying more on ankle impedance and reducing muscle activity compared to the other two tasks.

### B. Task-dependent contributions of impedance and muscle activations to net torque

We examined the contributions of ankle impedance and changes in muscle activation to the net torque at the ankle to determine the strategies that subjects employed in each task. Figure 4A shows the measured torque, the torques attributed to the impedance and muscle activation and the sum for a representative subject. Figure 4B shows the magnitude and the phase of the 0.5 Hz component of these torques for all subjects. For both the representative subject and the entire group, the torque due to impedance closely matched the measured torque in the +K task. In contrast, in the 0K task, the torque from impedance was a sinusoid of similar magnitude to the +K task, but not helpful as the 0K task required no sinusoidal torque component. Finally, the torque from impedance was a sinusoid out of phase with the measured torque in the -K task, and therefore counterproductive.Subjects used changes in muscle activation to compensate for counterproductive impedance torques. For all three tasks, the torque from ankle impedance was in phase with the movement (Fig. 4B). The timing of the impedance torque was therefore counterproductive in the 0K and -K tasks, requiring compensation through changes in muscle activation. The consequence was that subjects produced less torque from changes in muscle activation during the +K task and more torque due to ankle impedance (Δ = 4.9 %MVC, %95CI = 2.8–7.1, t_9_ = 5.3, p = 0.0005) (Fig. 4C). These contributions to the net ankle torque were more comparable in the 0K task (Δ = 1.8 % MVC, %95CI = 0.3–3.3, t_9_ = 2.8, p = 0.0207), while there was greater torque due to muscle activation in the -K Task (Δ = -12.3% MVC, %95CI = -14.2–-10.3, t_9_ = -17.5, p < 0.0001). Together, these results suggest that subjects relied more heavily on the impedance of the ankle to meet the task demands when impedance was consistent with the task requirements.

In all tasks, the torque due to impedance was dominated by the stiffness, resulting in as impedance torque that was in phase with the movement. The non-parametric IRFs quantifying ankle impedance were parameterized by second-order models with stiffness, viscosity and inertia (Figure 5). The torque due to stiffness (+K: 10.9 ± 1.5% MVC; 0K: 11.2 ± 2.6% MVC; -K: 7.9 ± 1.9% MVC, n = 10 for each task) was an order of magnitude greater than the torque due to viscosity torque (+K: 0.79 ± 0.20% MVC; 0K: 0.96 ± 0.28% MVC; -K: 1.06 ± 0.23% MVC) and two orders of magnitude greater than the torque due to inertia (0.16 ± 0.05% MVC in all tasks) in all three tasks.

**Figure 5.**
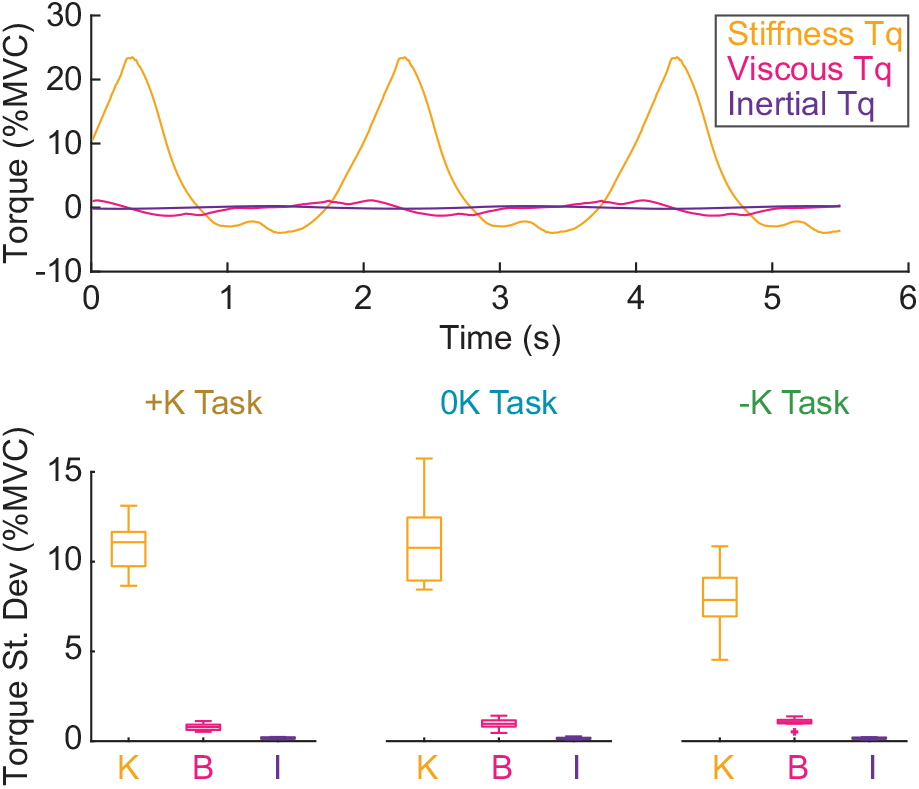
Stiffness was the dominant contributor to the impedance torque. A) The torque due to, stiffness (orange) viscosity (magenta) and inertia (purple) for a single task in a representative subject. B) The torque due to stiffness was approximately an order of magnitude greater than the torque due to viscosity and two orders of magnitude greater than the torque due to inertia for all tasks.

### C. Tasks that leverage limb impedance reduce the need for muscle activation

We compared the torque from muscle activation across the three tasks (Fig. 6A) as one measure of how each task influenced neural control. We found that the torque due to muscle activation was smallest in the +K task. The mean torque due to muscle activation per cycle (F_2,18_ = 11, p = 0.0008), an indication of average effort, and the standard deviation of the torque (F_2,18_ = 150, p < 0.0001), an indication of the complexity of neural control, varied between the different tasks. The mean torque from muscle activation was significantly lower in the +K task compared to the -K task (Δ = 7.5% MVC, %95CI = 2.9–12.1, p = 0.0017) and the 0K task (Δ = 7.2% MVC, %95CI = 2.6–11.9, p = 0.0023). The standard deviation was also smallest in the +K task compared to the -K task (Δ = 14.5% MVC, %95CI = 12.3–16.7, p < 0.0001) and the 0K task (Δ = 3.4% MVC, %95CI = 1.2–5.6, p = 0.0030). These results suggest that leveraging impedance in the +K task resulted in a physically easier and simpler task to perform.

**Figure 6.**
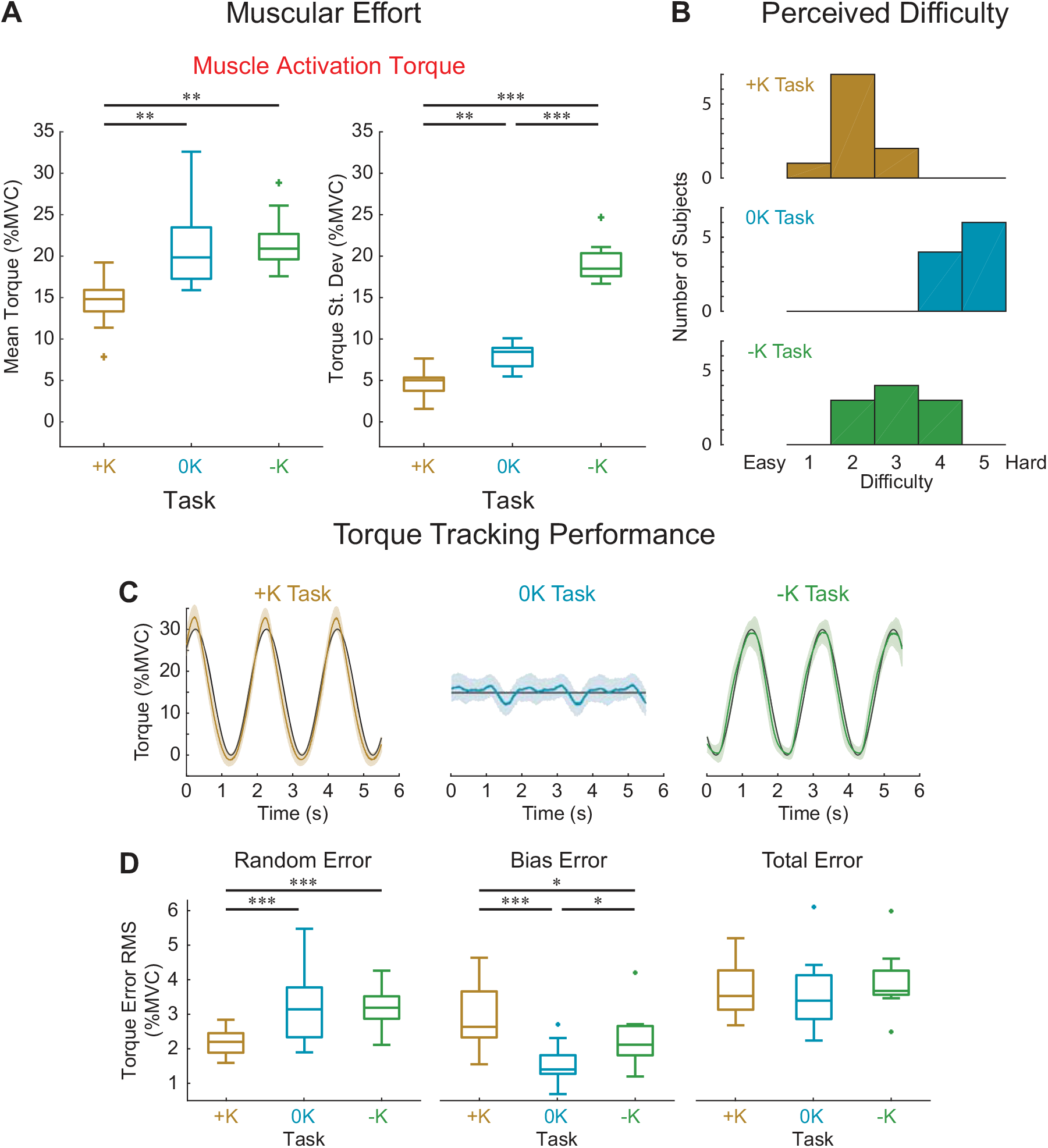
Leveraging impedance when feasible resulted in an easier and more consistent task performance. A) The +K task required less muscular effort to complete as subjects generated less torque due to muscle activation. This was seen both in the mean torque, indicative of the overall physical effort, as well as the standard deviation, indicative of a lesser need for precisely timed muscle activation that required mental focus. B) The +K task was perceived to be easier by the subjects, as rated on a 1–5 difficulty scale. C) Task performance was quantified by how well the measured torque matched the torque target in all three tasks. Representative data for one subject, showing the target (black), the average measured torque (colored lines) and standard deviation across all the cycles of the movement (shaded area). D) While the total error did not differ between the three tasks, the +K task did show less random error, indicating a more consistent task behavior when subjects could rely more heavily on impedance.

To rule out the possibility that co-contraction resulted in an increase in muscle activation with no increase in joint torque, we verified that the plantarflexor EMG activity was lower in the +K task. We found that the mean plantarflexor EMG was significantly lower in the +K task compared to the 0K task (Δ = 5.8% MVC, %95CI = 2.7–9.0, p = 0.0005) or the -K task (Δ = 4.4% MVC, %95CI = 1.2–7.5, p = 0.0064). This lesser amount of plantarflexor EMG in the +K task rules out the possibility that co-contraction of the plantarflexors and dorsiflexors resulted in low levels of torque due to muscle activation despite high levels of muscle activity.

### D. Tasks that can leverage limb impedance are perceived as easy to perform

We asked subjects to rate the difficulty of each task to determine how the different strategies we observed changed the perception of task difficulty. Difficulty was rated, from 1–5 on a continuous scale. These subjective measures were obtained from a new set of subjects so that previous exposure to the three torque-tracking tasks did not alter perceived difficulty. Eight of the 10 subjects perceived the +K task to be easiest, while 2 subjects found the -K task to be easiest. All subjects found the 0K task to be most difficult. Using the subjects’ reported perceived difficulty scores (Fig. 6B), we found that the there was a significant perceived difficulty between the three tasks (F_2,18_ = 38, p < 0.0001). The +K task was found to be significantly easier than both the –K (Δ = 1.0, %95CI = 0.2–1.7, p = 0.0091) and the 0K tasks (Δ = 2.5, %95CI = 1.7–3.2, p < 0.0001).

Not only did subjects perceive the +K task to be the easiest, their qualitative feedback was consistent with a task that required less effort. For example, subjects noted that it felt like they had to “push less”, possibly reflecting the decrease in torque due to muscle activation. Other subjects may have attributed the simplicity of the +K task to the decrease in cyclic muscle activation as they stated that they found the +K task easiest because all they had to do was “hold constant” or “hold their foot still”.

### E. Tasks that leverage limb impedance can be performed more consistently than others

We assessed the torque-tracking errors to evaluate how the different control strategies influenced performance in each task. Specifically, we quantified the total, random, and bias tracking errors for each subject (Figs. 6C&D). We saw no difference in the total tracking error (F_2,18_ = 1.63, p = 0.223). This similar performance may be due to fact that subjects were trained to achieve a certain level of proficiency in matching the torque. However, we did observe a differences in random error (F_2,18_ = 15.93, p = 0.0001). There was significantly less random error— and hence more consistent task performance—in the +K task compared to the -K (Δ root mean square error = 1.0% MVC, %95CI = 0.5–1.5, p = 0.0004) and 0K tasks (Δ = 1.0% MVC, %95CI = 0.5–1.5, p = 0.0003). The lower amount of random error is consistent with relying less on inherently noisy muscle activation (Carlton *et al*., 1985; Jones *et al*., 2002; Tracy *et al*., 2005). We also saw differences in bias error (F_2,18_ = 20.29, p < 0.0001), between the three tasks. There was greater bias error in the +K task compared to the -K (Δ = 0.7% MVC, %95CI = 0.1–1.2, p = 0.0131) and 0K tasks (Δ = 1.4% MVC, %95CI = 0.8–1.9, p < 0.0001). The bias error may be due to subjects adopting a simpler strategy. Since there was no penalty for not matching the task perfectly, and subjects needed to only achieve a level of proficiency similar to the other two tasks, subjects may have decided that this bias error was an acceptable trade off to perform the task simply and efficiently. It is important to note that in our study the movement was completely predictable, and may have aided subjects in producing muscle activation torques on time. Any small shifts in the timing of movement, as is normally seen in walking (Jordan *et al*., 2007), may have hindered subjects’ ability to correctly predict the timing of the muscle activation and hindered the performance of the strategies placing a greater reliance on muscle activation. Combined, these results demonstrate how relying on impedance in the +K task instead of timed muscle activation resulted in a more consistent and simpler task.

## DISCUSSION

The purpose of this study was to determine if leveraging the impedance of a limb to perform a task results in a simpler and more efficient behavior. We had subjects complete three torque-tracking tasks using the ankle, only one of which could be achieved via by leveraging their impedance (the +K task). We evaluated the control strategy used in each task by estimating the contributions of ankle impedance and muscle activation to the net torque that was produced. We found that subjects generated less torque from muscle activation in the +K task, indicative of less physical effort. Subjects also perceived the +K task to be easiest and were able to complete it more consistently than the other two tasks. Combined these results demonstrate that when subjects can leverage their joint impedance to complete a motor task, it results in a strategy that is easier and more consistent to perform.

The mechanical impedance of human limbs has been studied extensively in the context of maintaining stability during postural control (Hogan, 1984; Mussa-Ivaldi *et al*., 1985; De Serres & Milner, 1991; Trumbower *et al*., 2009; Krutky *et al*., 2013) and movement (Gomi & Kawato, 1996; Burdet *et al*., 2001; Franklin *et al*., 2007; Zenzeri *et al*., 2014). When stability is compromised by unexpected disturbances or the exertion of forces on the environment, we are able regulate impedance so as to complete the task at hand (Hogan, 1985a). Impedance regulation has also proven to be an important concept for robot control (Anderson & Spong, 1988; Vanderborght *et al*., 2013), as it can be used to generate stable motions and postures along with reliable and forceful contact with the environment. Impedance in robotics has been implemented both through software (Semini *et al*., 2015) and hardware (Pratt & Williamson, 1995; Vanderborght *et al*., 2013), analogous to the roles of muscle activation and joint impedance play in generating torques in our study respectively. Using a hardware based approach can be simpler as it reduces the complexity of the required control algorithms (Vanderborght *et al*., 2013). We believe that our experimental findings are the first to demonstrate that humans can also leverage the impedance of their limbs to simplify the control required to generate forceful interactions with the environment.

Many common tasks require joint motion-torque relationships that are consistent with the impedance of our limbs. Due to the ability of our central nervous system to precisely control muscle activation, humans have the ability to make their limbs and joints display a variety of motion-torque patterns or mechanical behaviors. One behavior that arises in many movements is a spring-like behavior for which the generated joint torques or limb forces act to oppose changes in length. Spring-like behaviors can be seen at the whole limb level (Blickhan, 1989; Ferris & Farley, 1997) and at the joint level (Farley & Morgenroth, 1999; Shamaei *et al*., 2013a) during human locomotion. For example, the ankle, knee, and hip exhibit spring-like behaviors in a variety of tasks including, walking (Shamaei *et al*., 2013b, a, c; Rouse *et al*., 2014), running (Stefanyshyn & Nigg, 1998; Arampatzis *et al*., 1999; Kuitunen *et al*., 2002), and hopping (Farley & Morgenroth, 1999). This spring-like behavior is also present at the muscle-tendon level during many phases of animal locomotion (Tu & Dickinson, 1996; Roberts *et al*., 1997; Biewener *et al*., 1998). A simple and efficient way to achieve this spring-like behavior would be to match the impedance of the joint to the behavioral demands so that the demands on changes in muscle activation are minimized. Our results clearly demonstrate that task-appropriate impedance simplifies neural control. An important complementary experiment would be to evaluate if impedance is actively regulated to simplify neural control.

We focused on ankle movements that have a frequency and amplitude relevant to locomotion and demonstrated that the torques generated within this operating regime are dominated by ankle stiffness. Stiffness, however, is only one component of the net ankle impedance. For many conditions, impedance can be characterized by a second-order system with inertial, viscous, and stiffness components. It would be interesting to determine if behaviors that are consistent with these higher order components of impedance also simplify neural control. Damping behaviors would be of particular interest since they are important for many movements that require stopping or deceleration. Damping is also essential for controlling movement when interacting with purely inertial loads (Lin & Rymer, 2001; Ludvig *et al*., 2008). For example, the ankle plays a substantial role in damping out energy of a falling body when landing from a jump (Gross & Nelson, 1988; Schmitz *et al*., 2007) but it is unclear if this behavior arises from the intrinsic mechanics of the ankle or changes in neural control. An approach similar to that used in our current study could answer this question and expand our understanding of the tasks that can be simplified by leveraging joint and limb impedance.

Our results were obtained using a novel method for decomposing the net torque about the ankle into components arising from impedance and from muscle activation. While impedance was measured during the cyclical tasks that we studied, the model used to estimate the torque from muscle activation was constructed from data measured during isometric contractions. It is well known that the EMG-torque relationship changes during movement, as this relationship is sensitive to the angle of the joint (Liu *et al*., 2013; Liu *et al*., 2015) and the velocity of movement (Wickiewicz *et al*., 1984). These effects may have introduced errors in our predictions of muscle activation torque during our movement conditions, but our results suggest that these errors were modest and consistent across all tested conditions. We saw an overall error standard deviation of 4.0% MVC. The majority of this error is likely due to our time-varying impedance model, as we saw similar error magnitudes in the passive trials in the absence of muscle activity.

## CONCLUSION

In summary, we developed a novel method that allowed us to separately estimate the contributions of ankle impedance and changes in neural control to the net ankle torque during large movements. Using this method, we determined that humans can leverage the impedance of the ankle to simplify neural control when that impedance is consistent with the motion-torque demands of the task. Leveraging impedance reduced the required muscular effort, leading to performance that was perceived to be easier and was completed more consistently. These results suggest that relying on impedance could be a simple and efficient way to complete many tasks that require spring-like motion-torque profiles from the joints within the human body.

## DATA AVAILABILITY

The authors will make the data available upon reasonable request via email to the corresponding author (D.L.).

## AUTHOR CONTRIBUTIONS

All authors contributed to the conceptual design of the study. D.L. and M.W.W. collected and analyzed the data. D.L. and E.J.P interpreted the results. D.L., M.W.W. and E.J.P. drafted the manuscript, while D.L. generated all the figures. All authors have approved the final version of the manuscript and have agreed to be accountable for all aspects of the work. The authors confirm that all listed authors qualify to be listed as authors, and that no one else qualifies to be an author of this manuscript.

